# ETV2 primes hematoendothelial gene enhancers prior to hematoendothelial fate commitment

**DOI:** 10.1101/2021.06.25.449981

**Authors:** Jeffrey D. Steimle, Chul Kim, Rangarajan D. Nadadur, Zhezhen Wang, Andrew D. Hoffmann, Erika Hanson, Junghun Kweon, Tanvi Sinha, Kyunghee Choi, Brian L. Black, John M. Cunningham, Kohta Ikegami, Ivan P. Moskowitz

## Abstract

The lineage-determining transcription factor ETV2 is necessary and sufficient for hematoendothelial fate commitment. We investigated how ETV2-driven gene regulatory networks promote hematoendothelial fate commitment. We resolved the sequential determination steps of hematoendothelial versus cardiac differentiation from mouse embryonic stem cells. *Etv2* was strongly induced and bound to the enhancers of hematoendothelial genes in a common cardiomyocyte-hematoendothelial mesoderm progenitor. However, only *Etv2* itself and *Tal1*, not other ETV2-bound genes, were induced. Despite ETV2 genomic binding and *Etv2* and *Tal1* expression, cardiomyogenic fate potential was maintained. A second wave of ETV2-bound target genes was up-regulated during the transition from the common cardiomyocyte-hematoendothelial progenitor to the committed hematoendothelial population. A third wave of ETV-bound genes were subsequently expressed in the committed hematoendothelial population for sub-lineage differentiation. The shift from ETV2 binding to productive transcription, not ETV2 binding to target gene enhancers, drove hematoendothelial fate commitment. This work identifies mechanistic phases of ETV2-dependent gene expression that distinguish hematoendothelial specification, commitment, and differentiation.

**HIGHLIGHTS:**

- The hematoendothelial master TF ETV2 is expressed in a multipotent mesoderm progenitor.
- ETV2 binds to target genes in the mesoderm progenitor without target gene activation.
- ETV2 binding in the mesoderm progenitor does not diminish alternative cardiac fate potential.
- ETV2-bound target genes are activated upon hematoendothelial fate commitment.

**GRAPHICAL ABSTRACT:** 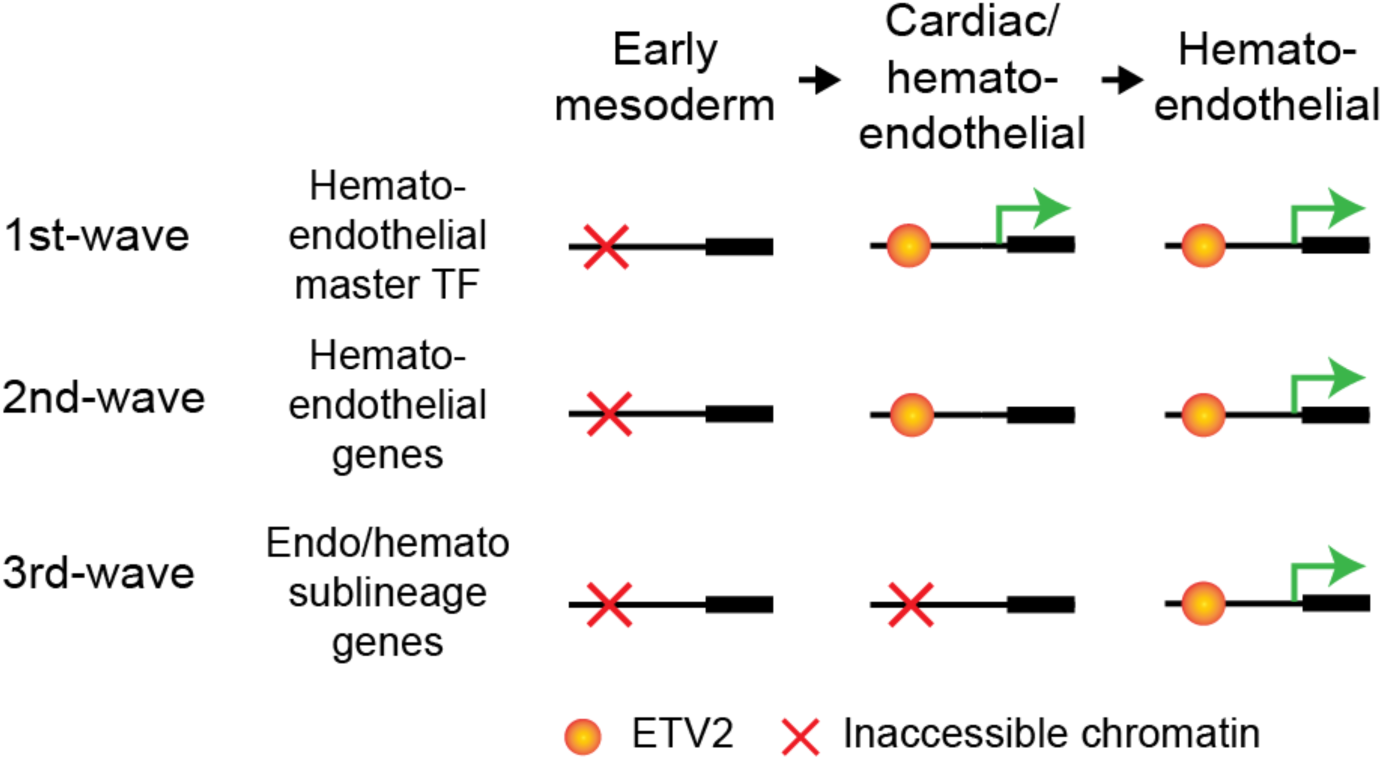

## INTRODUCTION

Lineage-determining transcription factors (TFs) commit cells to specific cell lineages during development (Spitz and Furlong, 2012). Lineage-determining TFs drive cell-type specific gene expression that triggers the cascade of cellular processes culminating in cell fate commitment. A prevailing model of cell lineage specification is that lineage-determining TFs drive terminal differentiation towards a specific lineage. An alternative model is that lineage-determining TFs “prime” progenitor cells toward a particular fate by chromatin binding, without activating lineage-specific gene expression programs within multipotent progenitors. Whether fate priming is the general feature of lineage-determining TFs remains unclear.

The Ets family transcription factor ETV2 is a lineage-determining TF for the vertebrate hematoendothelial lineage. Significant strides have been made to derive these fates *in vitro*, via biologically relevant intermediates using factors implicated *in vivo* in mouse genetic studies. Hematoendothelial fate specification in mouse embryonic stem cell (mESC) differentiation has been well characterized (Koyano-Nakagawa and Garry, 2017). Progenitor cells for the hematoendothelial lineage expresses Flk1 (VEGFR2), encoded by *Kdr*, the predominant receptor for vascular endothelial growth factors, VEGF (Choi et al., 1998; Ema et al., 2003; Kataoka et al., 1997; Park et al., 2004; Shalaby et al., 1995, 1997; Yamashita et al., 2000). VEGF-reception by Flk1 is the inductive cue for the hematoendothelial fate specification by transcriptionally inducing the hematoendothelial master transcription factor, *Etv2* (Kataoka et al., 2011; Rasmussen et al., 2012; Zhao and Choi, 2017). *Etv2* is necessary and sufficient to induce the hematoendothelial lineage (Kataoka et al., 2011; Koyano-Nakagawa et al., 2012; Lee et al., 2008; Park et al., 2004; Pick et al., 2007; Rasmussen et al., 2011, 2013) by directly binding and regulating genes that characterize the hematoendothelial lineage (De Val and Black, 2009; Garry, 2016; Koyano-Nakagawa and Garry, 2017; Liu et al., 2015).

Early mesoderm differentiation generates the vertebrate hematoendothelial lineage, a path for hematopoiesis and vasculogenesis, and the cardiac lineage, a path for heart development (De Val and Black, 2009). Interestingly, Flk1-positive cells give rise to both lineages *in-vivo* and *in-vitro*. The observation that cardiomyocytes and hematoendothelial lineages are both derived from early Flk1-positive mesoderm suggests that Flk1-positive mesoderm may represent a common progenitor for both lineages (Chen et al., 2006; Kattman et al., 2011; Motoike et al., 2003). How ETV2 induces hematoendothelial fate from the multipotent mesoderm progenitor remains unclear.

In this study, we examined fate intermediates originating from a single time point of mESC differentiation, thereby allowing us to investigate the molecular dynamics during a fate transition with a high temporal resolution. We investigated ETV2 binding and chromatin accessibility dynamics in mESC-derived mesoderm progenitors from these defined stages of hematoendothelial versus cardiogenic fate specification. Our data revealed an unexpected binding of ETV2 without transcriptional induction of hematoendothelial gene expression in multipotent mesoderm progenitors. Furthermore, ETV2 binding did not prevent the production of cardiomyogenic fate from the multipotent mesoderm progenitor population. Our results indicate that ETV2 is poised at the target genes in the multipotent progenitor population but does not act to drive hematoendothelial lineage gene expression until hematoendothelial fate specification.

## RESULTS

### Sequential derivation of PDGFRα and FLK1 expressing mesodermal progenitors from mouse embryonic stem cells

During early vertebrate development, cells expressing the PDGF-receptor PDGFRα and the VEGF receptor FLK1 are multipotent cardiovascular progenitors that can generate cardiomyocytes, vascular endothelium, and hematopoietic lineages (Choi et al., 1998; Ding et al., 2013; Ema et al., 2006; Kataoka et al., 1997; Yamashita et al., 2000). We aimed to identify the mechanisms underlying cardiovascular fate specification in the PDGFRα^+^ FLK1^+^ cell population. We delineated the mouse embryonic stem cell (mESC) differentiation into the PDGFRα^+^ FLK1^+^ population in order to isolate specific intermediates. We differentiated mESCs using a serum-free defined medium containing <u>A</u>ctivin A, <u>B</u>MP4, and <u>V</u>EGF (ABV regimen), previously described for mesoderm induction (Kattman et al., 2006, 2011). We isolated the PDGFRα^+^ FLK1^+^ (*PF*) population, the PDGFRα^+^ FLK1^−^ (*P*) population, and the PDGFRα^−^ FLK1^+^ (*F*) population at Day 4 of differentiation and examined their fate potentials in basal medium without ABV (**Fig. 1A**). The sorted *PF* population robustly differentiated into cardiac troponin T (cTnT)-positive beating cardiomyocytes (**Fig. 1B; Supplemental Movie 1**), consistent with the previous observations (Kattman et al., 2006, 2011). On the other hand, the sorted *P* and *F* populations did not differentiate into cardiomyocytes (**Fig. 1B**). The sorted *F* population robustly formed blood colonies, a feature characteristic of the hematoendothelial progenitor, as previously reported (Kattman et al., 2006, 2011) (**Fig. 1C & S1A**). The sorted *PF* and *P* populations did not form appreciable blood colonies (**Fig. 1C & S1A**). Therefore, the *PF* population appeared to define a cardiomyogenic progenitor, the *F* population a hematoendothelial population, and the *P* population lacked both cardiomyogenic and hematoendothelial potentials.

**Figure 1.**
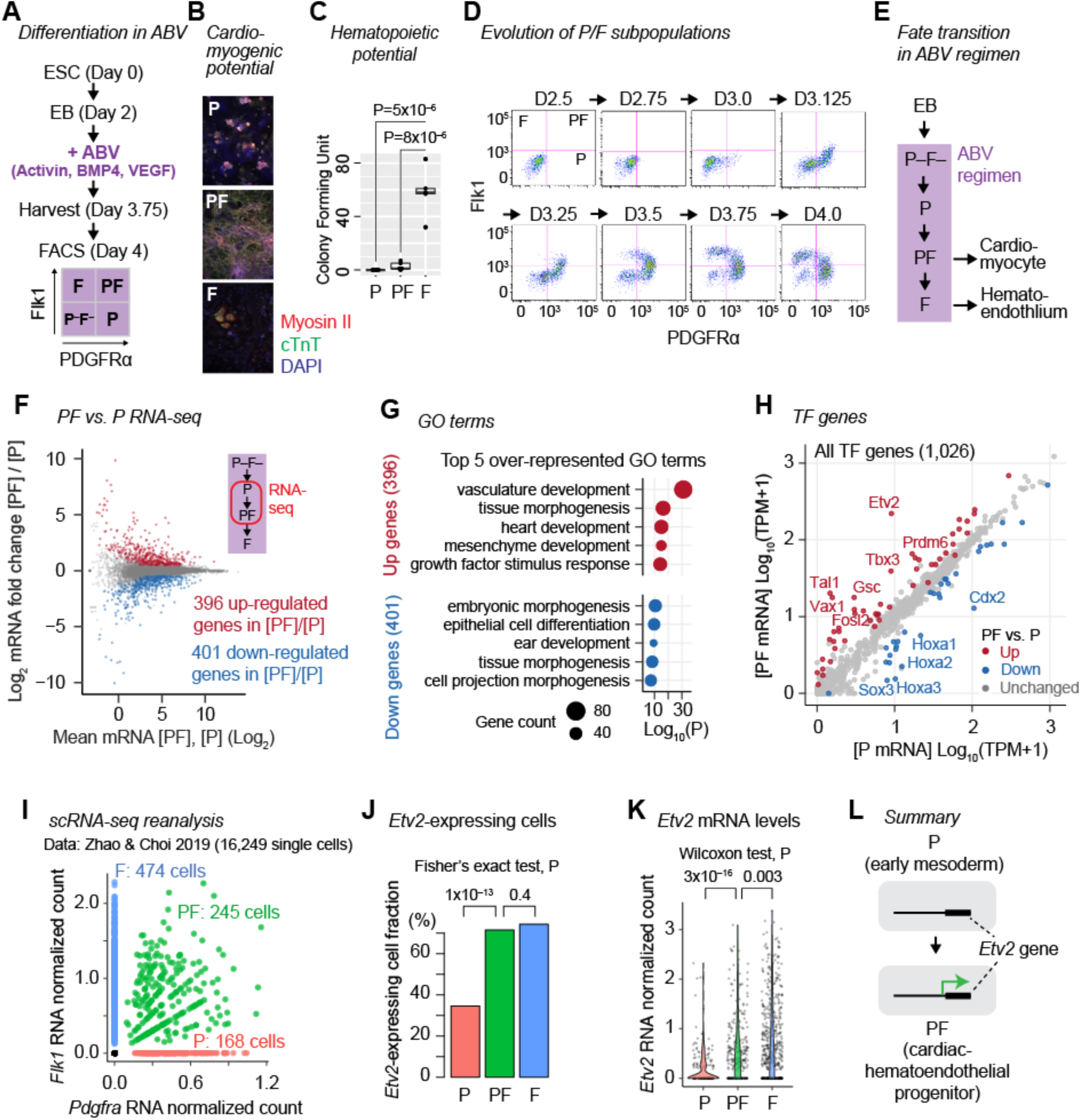
Hematoendothelial master TF *Etv2* is expressed in the *PF* cardiovascular progenitor. (A) mESC differentiation in the ABV regimen and isolation of PDGFRα/Flk1 subpopulations. (B) Immunofluorescence for cardiac Myosin II and cardiac troponin (cTnT) in the *P, PF*, *F* populations at 4 days after isolation. (C) Blood colony forming assays of the *P*, *PF*, *F* populations at 12 days after isolation. Results for individual blood subtypes are shown in **Fig. S1**. (D) Flow cytometry analysis for PDGFRα and Flk1 on cells harvested at the indicated time of differentiation. (E) The temporal relationship between the PDGFRα/Flk1 subpopulations. Purple box indicates the cell populations in the ABV regimen. (F) MA plot comparing RNA-seq TPMs (transcripts per million) of all genes in *PF* vs. *P* populations. (G) Top 5 most over-represented GO terms in the upregulated or downregulated genes in *PF* vs. *P*. (H) RNA-seq TPMs for all 1,026 transcription factor (TF) genes in the *P* (x-axis) and the *PF* (y-axis) populations. (I) *Pdgfra* and *Flk1* (*Kdr*) mRNA levels (normalized read counts) of 2,202 single cells (Zhao & Choi, 2019). *P*, cells with *Pdgfra* count > 0 and *Flk1* count = 0. *PF*, cells with *Pdgfra* count > 0 and *Flk1* count > 0. *F*, cells with *Pdgfra* count = 0 and *Flk1* count > 0. (J) Fraction of single-cell *P*, *PF*, or *F* cells expressing *Etv2* (*Etv2* read count > 0). (K) *Etv2* expression levels of single cells categorized as *P*, *PF*, or *F*. (L) Summary of Figure 1. Hematoendothelial master TF *Etv2* is strongly upregulated in the multipotent *PF* population.

To resolve the temporal relationship between the appearance of PDGFRα/FLK1 subpopulations, we analyzed PDGFRα and FLK1 expression every 3-6 hours from Day 2.5 to Day 4.0 of differentiation in the ABV regimen (**Fig. 1D; Fig. S1B**). EB-derived cells were initially PDGFRα^−^ FLK1^−^ (P–F–) at Day 2.5. The *P* population emerged at Day 3.0, followed by the *PF* population at Day 3.125, and finally the *F* population at Day 3.5. To further clarify the order of differentiation, we isolated PDGFRα/FLK1 subpopulations at Day 3.125 and individually cultured in the ABV regimen for 24 hours (**Fig. S1C**). The isolated P–F– cells gave rise to the *P* (49.5%) and *PF* populations (42.0%). The isolated *P* cells gave rise to *PF* (36.1%) and *F* (60.4%) populations. The isolated *PF* cells gave rise to the *F* population (60.6%) or remained *PF* (34.2%), but did not produce any appreciable *P* population (3.5%). This time series defined the temporal order of progenitor appearance as *P*, then *PF*, then *F*. This defined the *PF* population as a common cardiac/hematoendothelial progenitor that could differentiate into either cardiomyocytes upon isolation from the ABV regimen or into the *F* hematoendothelial population if retained in the ABV regimen (**Fig. 1E**).

### *Etv2* is expressed in the PDGFRα^+^ FLK1^+^ multipotent progenitor population

Characterization of fate intermediates originating from the single time point of mESC differentiation provided us with a unique opportunity to investigate the dynamics of chromatin accessibility, TF binding and gene expression during sequential phases of cell fate commitment. We investigated the molecular differences between the *P* population and *PF* population to identify the mechanisms for the cardiovascular fate acquisition in the *PF* population. We examined the transcriptomes of the *P* and *PF* populations sorted at Day 4 of differentiation in the ABV regimen by RNA-seq. We identified 396 upregulated and 401 downregulated genes in the *PF* population relative to the *P* population (**Fig. 1F; Table S1**). The upregulated genes were most strongly overrepresented for the gene ontology (GO) term “vasculature development” (P=10^−41^) reminiscent of the hematoendothelial fate (**Fig. 1G**), although the *PF* population did not present hematoendothelial potential (**Fig. 1C**). Consistent with this observation, *Etv2*, the master transcription factor (TF) orchestrating the hematoendothelial fate commitment, was among the most strongly upregulated TF genes in the *PF* population relative to the *P* population (26.4-fold upregulation; **Fig. 1H**). Furthermore, the *Etv2* mRNA level in the *PF* population was among the highest (98 percentile) in all 1,026 TF genes in the genome in the *PF* population (**Fig. 1H**). *PF* upregulated genes were also overrepresented for the GO term “heart development” (P=10^−27^) (**Fig. 1G**), including TFs implicated in cardiac development such as *Gata4*, *Hand2*, *Tbx3* and *Tbx20* (**Fig. 1H**) and consistently with the cardiomyogenic potential of the *PF* population (**Fig. 1B**).

The strong upregulation of *Etv2* in the *PF* multipotent population was unexpected given the known role of *Etv2* as a lineage-determining TF for hematoendothelial differentiation (Koyano-Nakagawa and Garry, 2017). To independently verify the elevated expression of *Etv2* in the *PF* population, we analyzed public single-cell transcriptomic data for mESCs undergoing spontaneous differentiation to various lineages in a serum-containing medium (Zhao and Choi, 2019), a condition different from the directed mesoderm differentiation offered by the ABV regimen. We computationally binned the single cells into the *P* (168 cells), *PF* (245 cells), and *F* (474 cells) populations based on the presence or absence of the mRNA expression of *Pdgfra* (encoding PDGFRα) and *Kdr* (encoding FLK1) (**Fig. 1I**). We observed that the *Etv2*-expressing cell fraction increased from 35% among the single-cell *P* population to 71% among the single-cell *PF* population (*P* vs. *PF*, Fisher exact test P=1×10^−13^) and then plateaued at 74% among the single-cell *F* population (*PF* vs. *F*, Fisher exact test P=0.4) (**Fig. 1J**). Concomitantly, the *Etv2* mRNA level in individual cells increased strongly in the single-cell *PF* population relative to the single-cell *P* population (Wilcoxon test P=3×10^−16^) and increased modestly in the single-cell *F* population relative to the single-cell *PF* population (Wilcoxon test P=0.003) (**Fig. 1K**). These results confirmed the unexpected observation that the hematoendothelial master TF *Etv2* was strongly expressed in the *PF* multipotent cardiovascular/hematoendothelial progenitor population (**Fig. 1L**).

### ETV2-binding sites gain chromatin accessibility in the PDGFRα^+^ FLK1^+^ cells

The strong upregulation of *Etv2* in the *PF* population led us to hypothesize that ETV2 might bind to chromatin in the *PF* population, potentiating the *PF* population for hematoendothelial fate. To define genomic differences between the *P* and *PF* populations, we examined accessible chromatin by ATAC-seq in these two populations. We identified 163,723 accessible sites detected in either *P* or *PF* population or both. Of these accessible chromatin sites, 5,847 sites (3.6%) gained accessibility whereas 2,511 sites (1.5%) lost accessibility from *P* to *PF*, respectively (**Fig. 2A; Table S2**). The Ets TF family DNA binding motif was the most strongly overrepresented motif in the gained accessibility sites (45% of the gained sites). Specifically, the ETV6 motif, essentially identical to the ETV2 motif (Liu et al., 2015; Neveu et al., 2018), was the most significantly overrepresented TF motif within the Ets family (**Fig. 2B**). In addition, TF family motifs for GATA (highest score, GATA4), bHLH (highest score, TAL1), T-box (highest score, TBX21), and Sox (highest score, TCF7) were overrepresented in the gained accessibility sites (**Fig. 2B**). The lost accessibility sites were overrepresented for the motifs of pluripotency TFs such as Sox (highest score TF, SOX2), homeodomain (highest score TF, NANOG), and the homodomain-POU family (highest score TF, POU5F1) (**Fig. 2B**). Thus, in the transition from *P* to *PF*, accessible sites utilized by pluripotency TFs were replaced by accessible sites utilized by TFs for both cardiac and hematoendothelial lineage specification.

**Figure 2.**
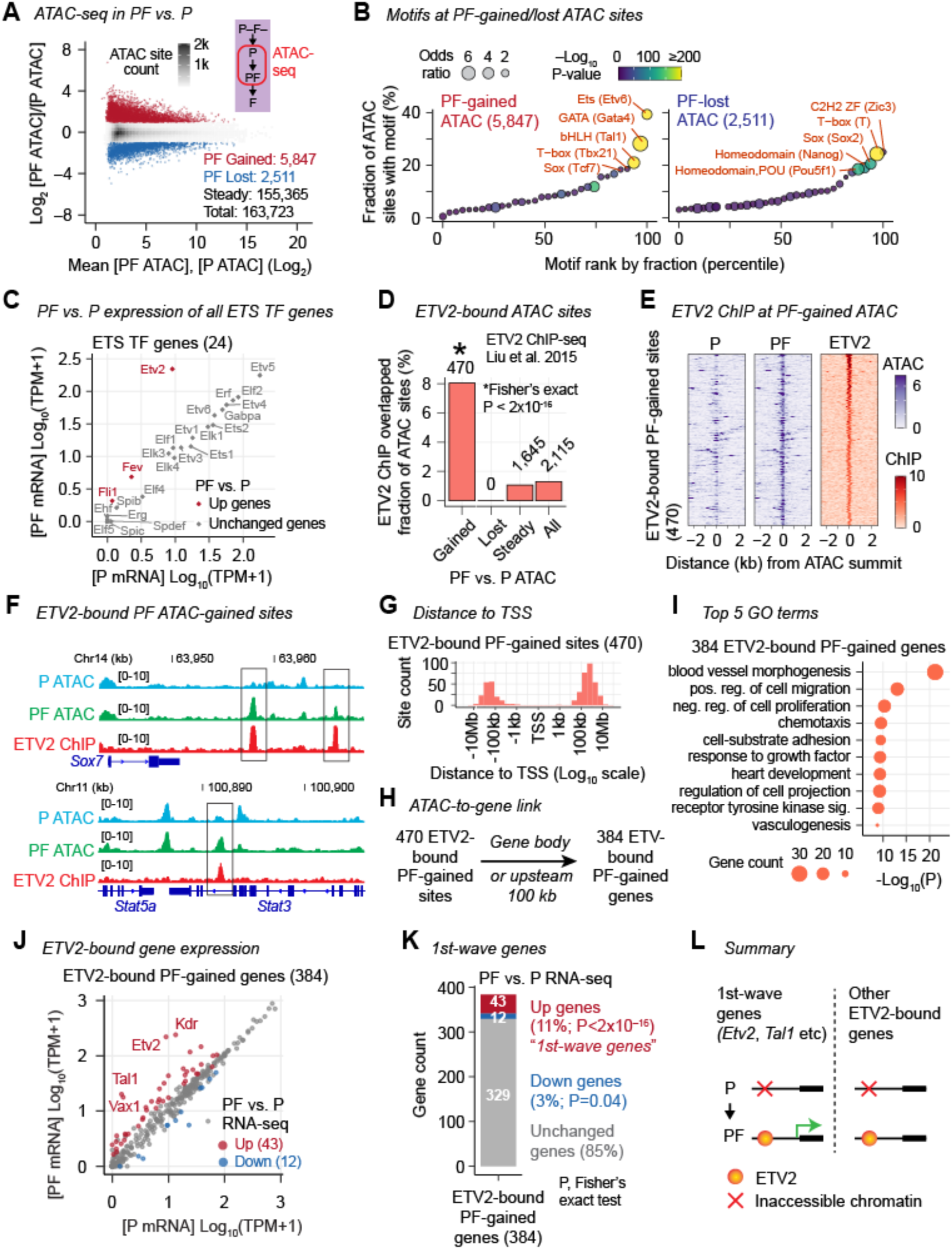
ETV2-binding sites gain chromatin accessibility in the *PF* population. (A) MA plot comparing ATAC-seq-derived chromatin accessibility at the 163,723 union ATAC sites in the *PF* population vs. the *P* population. (B) Transcription factor (TF) motifs over-represented (odds ratio > 1) within gained ATAC sites (left) or lost ATAC sites in *PF* vs. *P*. TFs are grouped by TF families, and the scores for the most over-represented motif within each family are shown (see **Methods**). (C) RNA-seq TPMs for all 24 ETS family TF genes in the *P* (x-axis) and the *PF* (y-axis) populations. (D) Fraction of *PF*-vs-*P* differentially accessible sites intersecting ETV2-binding sites. ETV2-binding sites are defined by ChIP-seq in FLK1-positive cells (Liu et al. 2015). (E) ATAC-seq and ETV2 ChIP-seq fold enrichment signals at the 470 *PF*-gained ATAC sites overlapping ETV2-binding sites (“ETV2-bound *PF*-gained sites”). (F) ATAC and ETV2 ChIP-seq fold enrichment signal tracks showing representative ETV2-bound *PF*-gained sites. (G) Histogram for the distance between the 470 ETV2-bound *PF*-gained sites and the closest transcription start site (TSS). (H) Algorithm to link genes to the 470 ETV2-bound *PF*-gained sites. The 384 genes linked are termed “ETV2-bound *PF*-gained genes.” Additional analyses on ETV2-bound PF-gained genes are shown in **Fig. S2**. (I) Top 5 most over-represented GO terms among the 384 ETV2-bound *PF* gained genes. (J) RNA-seq TPMs for the 384 ETV2-bound *PF*-gained genes in the *P* (x-axis) and the *PF* (y-axis) populations. (K) Number of upregulated, downregulated, or unchanged genes in *PF* vs. *P* among the 384 ETV2-bound *PF*-gained genes. The upregulated ETV2-bound *PF*-gained genes are termed the “first-wave genes.” (L) Summary of Figure 2. The first-wave genes are upregulated and bound by ETV2, as evidenced by the accessibility gain at ETV2-binding sites, in *PF*. Other genes are not upregulated in *PF* despite ETV2 binding in *PF*.

We asked whether ETV2 binding might be responsible for the accessibility gains at the ETS-family motif sites. We examined transcriptional states of all ETS family transcription factors in the genome and observed that three ETS TFs, *Etv2*, *Fli1*, and *Fev*, were significantly upregulated in *PF* relative to *P*. *Etv2* was the most upregulated (26-fold, FDR=3×10^−55^) and highest expressed (transcripts per million (TPM) = 218) TF in *PF* among all 24 ETS family TFs (**Fig. 2C**). *Fli1* and *Fev* were only modestly upregulated and expressed very weakly in *PF* (*Fli1*, 4.3-fold, FDR=1×10^−3^, TPM=1.1; *Fev*, 2.8-fold, FDR=0.02, TPM=3.8) (**Fig. 2C**). These data supported the hypothesis that ETV2 binding resulted in accessibility gains at ETS-family motif sites in the *PF* population.

To determine whether ETV2 bound to the gained accessibility sites in the *PF* population, we analyzed public ChIP-seq datasets for ETV2 performed in mESC-derived FLK1^+^ cells, which would include both the *PF* and *F* populations (Liu et al., 2015). This defined 3,868 ETV2-binding sites in the FLK1^+^ population (**Methods; Table S3**). These ETV2-binding sites were strongly overrepresented among the *PF*-gained accessibility sites (Fisher’s exact test P<2×10^−16^) and overlapped with 470 *PF*-gained-accessibility sites (8% of all gained accessibility sites; **Fig. 2D, E**). None of the ETV2-binding sites overlapped the lost accessibility sites (**Fig. 2D**). We termed the 470 *PF* gained-accessibility sites bound by ETV2 “ETV2-bound *PF*-gained sites”. These ETV2-bound *PF*-gained sites were found in genes including transcription factors *Sox7* and *Stat3* (**Fig. 2F**). Almost all of the ETV2-bound *PF*-gained sites were located outside of the TSS proximal (+/– 2 kb) regions (457, 97%) (**Fig. 2G**). We linked 384 genes, located within the gene bodies or 100-kb upstream regions to 402 of the 470 ETV2-bound *PF*-gained sites (“ETV2-bound *PF*-gained genes”) (**Fig. 2H; Table S1; Methods**). The vast majority (367, 96%) of the ETV2-bound *PF*-gained genes had a single ETV2-bound *PF*-gained site within the search space (**Fig. S2A**), and almost all (372, 97%) ETV2-bound *PF*-gained genes were linked only to gene-distal ETV2-bound *PF*-gained sites (outside +/–2 kb of TSS) (**Fig. S2B**). ETV2-bound *PF*-gained genes were strongly overrepresented for the GO term “blood vessel morphogenesis” and included known ETV2-target genes including *Etv2*, *Tal1*, *Kdr*, *Sox7*, and *Fli1* (**Fig. 2I; Table S1**). These results suggested that ETV2 bound to a subset of distal regulatory elements of the hematoendothelial specification genes in the *PF* cardiac/hematoendothelial multipotent population, prior to fate commitment.

### A small subset of ETV2-bound genes are upregulated in the PDGFRα^+^ FLK1^+^ population

ETV2 possesses a strong transcriptional activation domain (De Haro and Janknecht, 2002). We therefore asked whether the chromatin accessibility gains at ETV2-binding sites accompanied transcriptional upregulation of the associated genes in the *PF* population. Of the 384 ETV2-bound *PF*-gained genes, only 43 genes were upregulated in the *PF* population relative to the *P* population (11%; Fisher’s exact test P < 2×10^−16^) (**Fig. 2J, K**). We term these upregulated ETV2-bound *PF*-gained genes the “first-wave” genes (**Table S1**). The first-wave genes included *Etv2* itself and *Tal1*, a known immediate target gene transcriptionally activated by ETV2 (Kataoka et al., 2011; Liu et al., 2015; Wareing et al., 2012; Zhao and Choi, 2017). *Etv2* and *Tal1* constitute the transcriptional kernel for the hematoendothelial specification (Koyano-Nakagawa and Garry, 2017; Zhao et al., 2017). Twelve ETV2-bound *PF*-gained genes were downregulated in the *PF* population relative to the *P* population (3%; Fisher’s exact test P = 0.04) (**Fig. 2J, K**). The remaining 85% of the ETV2-bound *PF*-gained genes were not upregulated or downregulated. These results suggested that ETV2 binding to hematoendothelial genes did not accompany transcriptional upregulation in the multipotent *PF* population at the vast majority of the bound genes, except a small subset of genes including *Etv2* and *Tal1*, despite the known activator function of ETV2 (**Fig. 2L**).

### VEGF induces *Etv2* in the PDGFRα^+^ FLK1^+^ population

To define the direct consequence of ETV2 binding in the *PF* population, we investigated pathways that lead to *Etv2* expression and chromatin binding in the *PF* population. VEGF signaling through FLK1 is known to induce *Etv2* (Kataoka et al., 2011; Koyano-Nakagawa et al., 2015; Rasmussen et al., 2012; Zhao and Choi, 2017), suggesting that VEGF in the AB<u>V</u> regimen may induce *Etv2* in the *PF* population. To test this hypothesis, we characterized the effect of VEGF in the generation and gene expression of the *PF* population. Differentiation with VEGF (ABV regimen) or without VEGF (AB regimen) generated the *PF* population with comparable efficiency (**Fig. 3A**). Likewise, the FLK1 tyrosine kinase inhibitor ZD6474, which blocks VEGF signaling (Hennequin et al., 2002; Ryan and Wedge, 2005), did not alter the efficient generation of the *PF* population using the ABV regimen (**Fig. 3B**). Instead, the FLK1 inhibitor inhibited the subsequent generation of the *F* population from the *PF* population (**Fig. 3B**), consistent with the known requirement of VEGF for this transition (**Fig. 3B**) (Kennedy et al., 1997; Shalaby et al., 1995, 1997; Yamashita et al., 2000; Zhao and Choi, 2017). Thus, VEGF was dispensable for generating the *PF* population, but required for the transition from the *PF* population to the *F* population. To test whether VEGF induced *Etv2* in the *PF* population, we quantified *Etv2* mRNA in the *PF* population derived with or without VEGF. The *Etv2* mRNA level was 10-fold higher in *PF* derived with VEGF than in *PF* derived without VEGF (*t*-test P=0.004; **Fig. 3C**), indicating that VEGF induced *Etv2* in the *PF* population. Together, these observations indicated that *Etv2* is strongly induced in the transition from *P* to *PF* by VEGF but is itself dispensable for this transition.

**Figure 3.**
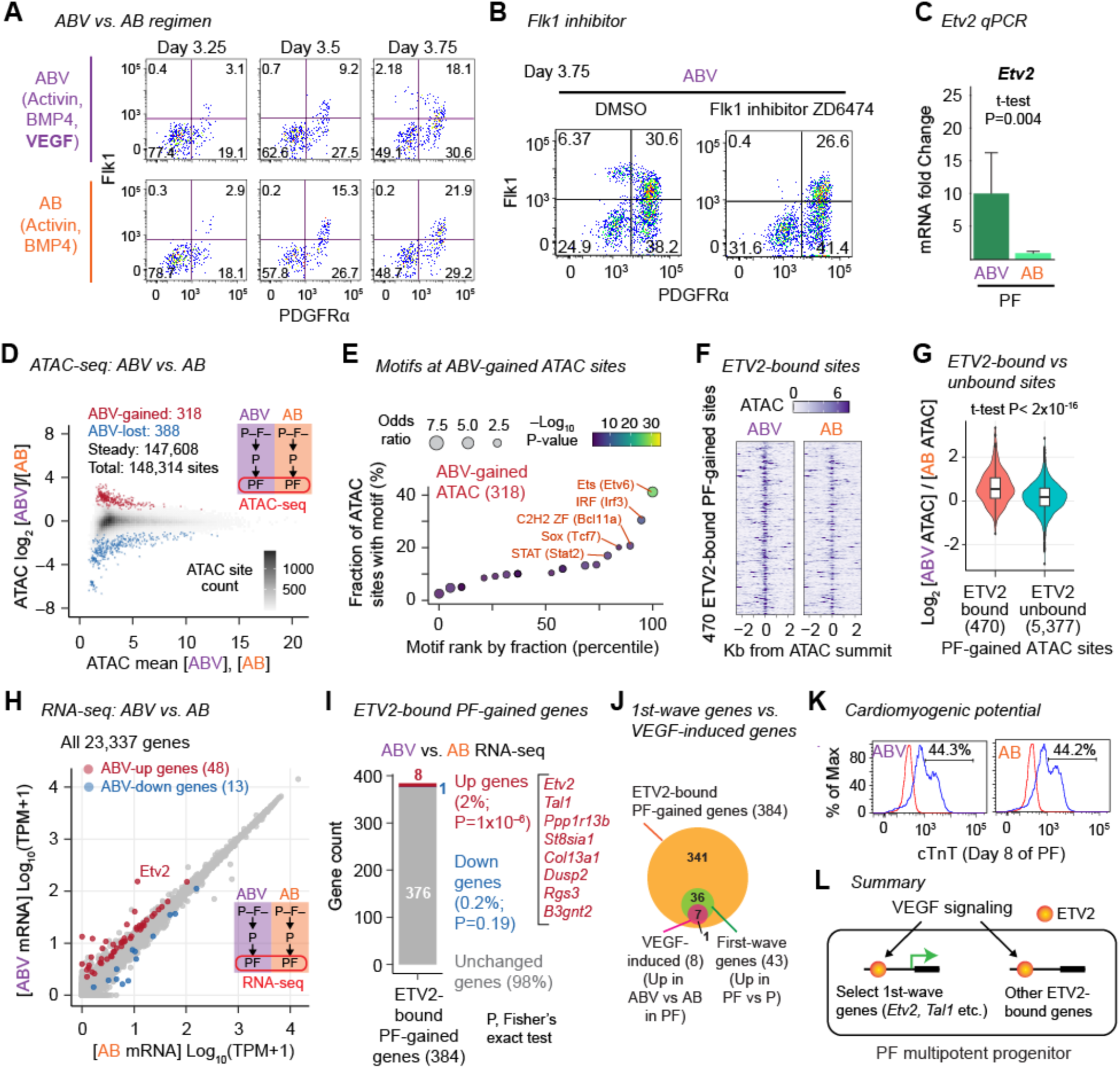
VEGF signaling drives ETV2 chromatin binding in the *PF* population. (A) Flow cytometry analysis for PDGFRα and FLK1 on cells harvested at the indicated time of differentiation in the ABV (top) or AB (bottom) regimen. (B) Flow cytometry analysis for PDGFRα and FLK1 on cells harvested at D3.75 of differentiation in the ABV regimen in the presence (right) or absence (left) of the FLK1 inhibitor ZD6474. ZD6474 was treated from day 2 to day 3.75 of differentiation. (C) RT-PCR analysis quantifying RNA levels of *Etv2* in the *PF* population derived with ABV or AB. The RNA levels are normalized to *Gapdh*. (D) MA plot comparing ATAC-seq-derived chromatin accessibility at the 148,314 union ATAC sites in the *PF* population derived with ABV vs. the *PF* population derived with AB. (E) TF motifs over-represented within the 314 gained ATAC sites in *PF* derived with ABV. TF motifs over-represented within the lost ATAC sites are shown in **Fig. S3**. (F) ATAC-seq fold enrichment signals at the 470 ETV2-bound *PF*-gained sites. (G) Log_2_ fold difference of ATAC-seq fold enrichment signals between *PF* derived with ABV and *PF* derived with AB at the 470 ETV2-bound *PF*-gained accessibility sites or the 5,377 ETV2-unbound *PF*-gained accessibility sites. (H) RNA-seq TPMs for all genes in *PF* derived with AB (x-axis) and *PF* derived with ABV (y-axis). (I) Number of upregulated, downregulated, or unchanged genes in *PF* derived with ABV vs. *PF* derived with AB among the 384 ETV2-bound *PF*-gained genes. The gene names of the 8 upregulated genes are indicated. (J) Relationship between the 43 first-wave genes (ETV2-bound *PF*-gained genes upregulated in *PF* relative to *P*) and the 8 VEGF-induced genes (ETV2-bound *PF*-gained genes upregulated in ABV relative to AV) among the 384 ETV2-bound *PF*-gained genes. (K) Flow cytometry analysis of the *PF* cells derived with ABV or AB, stained with the anti-cTnT antibody. (L) Summary of Figure 3. In the *PF* population, VEGF signaling induces ETV2 chromatin binding, and only a small fraction of ETV2-bound genes are induced in response to VEGF reception.

We predicted that chromatin accessibility at ETV2 binding sites in the *PF* population would depend on VEGF. To test this prediction, we profiled the chromatin accessibility in the *PF* population derived with VEGF (ABV regimen) or without VEGF (AB regimen), prepared in parallel by ATAC-seq. To our surprise, accessibility differences were detected at only 706 sites, with 318 sites showing higher accessibility and 388 sites showing lower accessibility in *PF* derived with VEGF versus *PF* derived without VEGF (**Fig. 3D; Fig. S3A; Table S4**). Nonetheless, the 318 sites with higher accessibility in *PF* derived with VEGF were strongly over-represented for Ets binding motif (**Fig. 3E**). Consistently, the 470 ETV2-bound *PF*-gained sites exhibited higher accessibility in *PF* derived with VEGF compared with *PF* derived without VEGF (**Fig. 3F**). Furthermore, the 470 ETV2-bound *PF*-gained sites showed stronger accessibility gains in *PF* derived with VEGF versus *PF* derived without VEGF, compared to the 5,377 *PF*-gained accessibility sites without ETV2 binding (*t*-test P< 2×10^−16^; **Fig. 3G**). These results supported the hypothesis that VEGF-dependent induction of *Etv2* resulted in ETV2 binding in the *PF* population, but were not required for the generation of the *PF* population itself.

### Very few ETV2-bound genes are activated directly by ETV2 in the PDGFRα^+^ FLK1^+^ population

We posited that the VEGF-dependency of *Etv2* expression and ETV2 binding in the *PF* population would allow us to identify the direct transcriptional targets of ETV2 in the *PF* population by manipulating VEGF. We profiled the transcriptomes of the *PF* population derived with VEGF (ABV regimen) or without VEGF (AB regimen) by RNA-seq. The transcriptomes of the *PF* population with VEGF and *PF* without VEGF were strikingly similar, with only 61 differentially expressed genes between the populations (**Fig. 3H**). These 61 genes consisted of 48 upregulated genes and 13 downregulated genes in *PF* with VEGF compared with *PF* without VEGF (**Table S1**). Strikingly, the most strongly upregulated genes, with greater than 10-fold upregulation, were *Etv2* and *Tal1* (**Fig. 3H**). This indicated that few ETV2 target genes are activated directly by ETV2 in the *PF* population generated with VEGF, despite strong expression of *Etv2* itself.

We then examined the expression change of the 384 ETV2-bound *PF*-gained genes. Of the 384 ETV2-bound *PF*-gained genes, only 8 genes showed higher expression in *PF* derived with VEGF compared with *PF* derived without VEGF (2.1%; Fisher’s exact test P=1×10^−6^) (**Fig. 3I**). These 8 ETV2-bound *PF*-gained, VEGF-induced genes were *Etv2*, *Tal1*, *Ppp1r13b*, *St8sia1*, *Col13a1*, *Dusp2*, *Rgs3*, and *B3gnt2* (**Fig. 3I; Table S1**). Among these genes, *Etv2*, *Tal1*, *Ppp1r13b*, and *Dusp2* are known transcriptional target genes of ETV2 (Gomez et al., 2009; Koyano-Nakagawa et al., 2015; Liu et al., 2015). The remaining 376 (98%) ETV2-bound *PF*-gained genes were not upregulated by VEGF in the *PF* population (**Fig. 3I**). All VEGF-induced ETV2-bound *PF*-gained genes, except *Rgs3*, were among the first-wave ETV2-bound genes upregulated in *PF* relative to *P*, as expected (**Fig. 3J**). The first-wave genes not induced by VEGF included *Kdr*, *Sox7*, *Fli1*, suggesting that these ETV2-bound genes were upregulated in the *PF* population by ETV2-independent mechanisms (**Fig. 3J**). Thus, ETV2 directly activated only the most upstream transcriptional kernel for the hematoendothelial specification, consisting of *Etv2* itself and the immediate ETV2 target *Tal1*, but remained dormant at the vast majority of ETV2-bound and accessibility-gained hematoendothelial genes in the *PF* population.

### ETV2 binding does not impede cardiomyogenic fate potential of the PDGFRα^+^ FLK1^+^ population

We tested whether ETV2-binding to the hematoendothelial genes and expression of the first-wave ETV2-bound genes in the *PF* population influenced the alternative cardiomyogenic fate potential of the *PF* population. We examined the cardiomyogenic fate potential of the *PF* population derived with VEGF (ABV regimen) or without VEGF (AB regimen). The *PF* population derived with and without VEGF exhibited comparable differentiation potential to cTnT-positive cardiomyocytes (44.3% with VEGF vs. 44.2% without VEGF) (**Fig. 3K**). Collectively, VEGF-induced ETV2 chromatin binding to hematoendothelial genes and activation of the first-wave ETV2-bound genes did not impede the alternative cardiomyogenic potential of the *PF* population (**Fig. 3L**).

### Genes that gained ETV2 binding in the PDGFRα^+^ FLK1^+^ population are extensively upregulated in the PDGFRα^−^ FLK1^+^ population

We hypothesized that ETV2-bound genes in the *PF* population might be primed for activation in the *F* population. To determine whether the ETV2-bound *PF*-gained genes would be activated in the *F* population, we profiled the transcriptome of the *F* population and compared it with the transcriptome of the *PF* population. We observed extensive gene expression differences, with 1,412 genes upregulated and 1,331 genes downregulated in *F* compared with *PF* (**Fig. 4A; Fig. S4A; Table S1**). The most overrepresented GO term among the 1,412 upregulated genes was “blood vessel development” consistent with the hematoendothelial fate of the *F* population (**Fig. S4A**). The overrepresented GO terms among the 1,331 downregulated genes included “heart development,” consistent with the loss of the cardiomyocyte differentiation potential during the transition from the *PF* to the *F* population (**Fig. S4B**). We then compared *PF* versus *F* differentially expressed genes with the 384 ETV2-bound *PF*-gained genes (**Fig. 4B**). The *F* upregulated genes were highly overrepresented among the ETV2-bound *PF*-gained genes (123 of 384 genes, 32%; Fisher’s exact test, P<2×10^−16^) but not among the downregulated genes (31 of 384 genes, 8%; P=0.06) (**Fig. 4C**). There were 123 ETV2-bound *PF*-gained genes upregulated in the *F* population, which we termed the “second-wave” ETV2-bound genes (**Fig. 4C; Table S1**). The second-wave genes were overrepresented for the GO term “blood vessel morphogenesis” (P=10^−12^), as expected (**Fig. 4D**). The second-wave ETV2-bound genes included 27 of the 43 first-wave ETV2-bound genes (63%) and all 8 VEGF-induced genes in *PF* (**Fig. 4E**), indicating that a majority of first-wave genes were further upregulated in the transition from *PF* to *F*. The second-wave genes included TFs known to play critical roles in hematoendothelial fate specification (*Etv2*, *Tal1*, *Fli1*) (Koyano-Nakagawa and Garry, 2017) or endothelial specification (*Sox7*, *Sox18*) (Yao et al., 2019) or repressing the hematopoietic expression program (*Klf7 and Etv6*) (Hock and Shimamura, 2017; Schuettpelz et al., 2012) (**Table S1**). In addition, strong overrepresentation of “regulation of Notch signaling pathway” (P=10^−8^), a signaling pathway required for hematoendothelial specification, was observed among the second-wave ETV2-bound *PF*-gained genes (Jang et al., 2015; Marcelo et al., 2013) (**Fig. 4D**). These results suggested that the second-wave genes were primed by ETV2 in the *PF* multipotent population for activation in the *F* hematoendothelial population.

**Figure 4.**
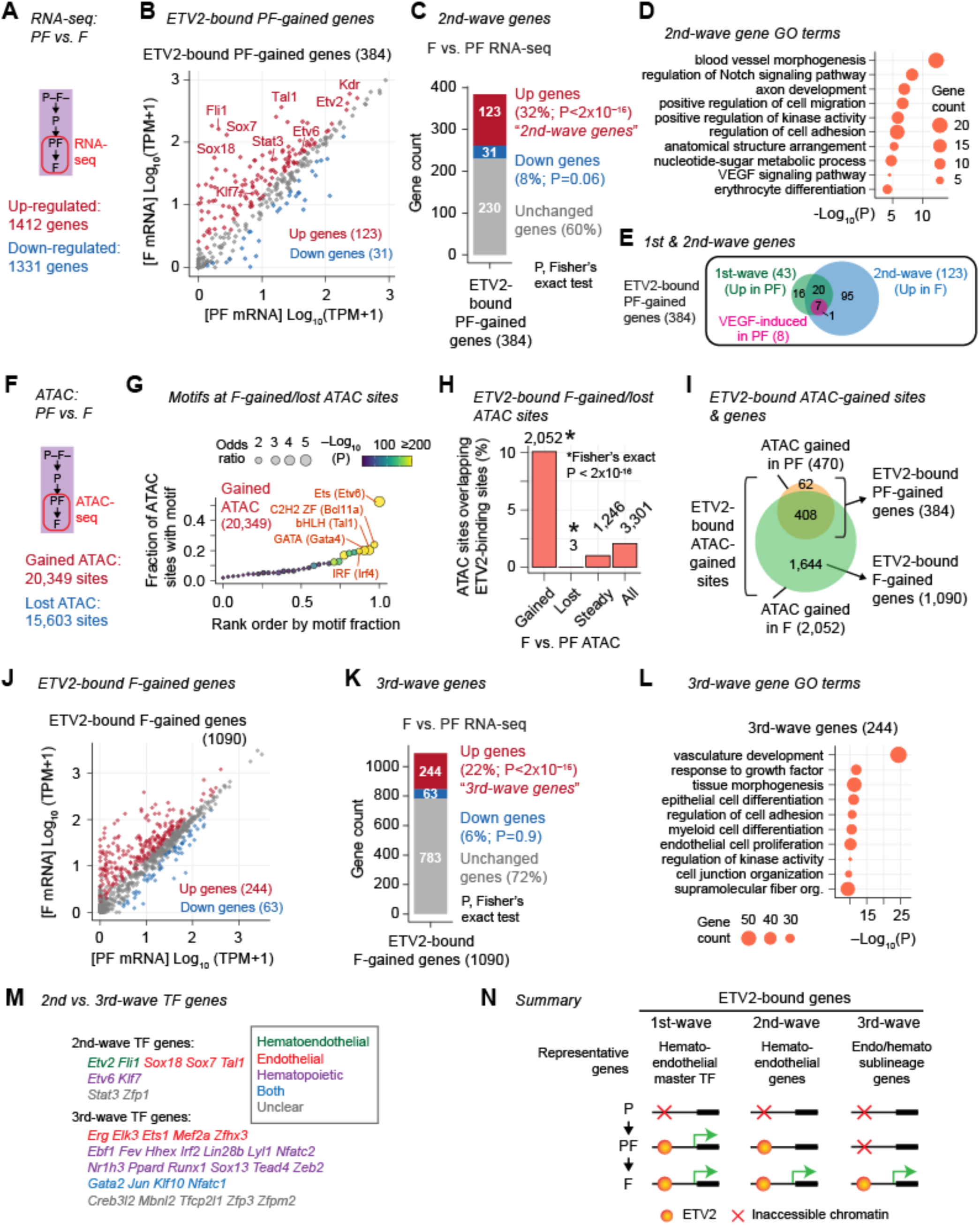
ETV2-bound genes are activated in the F hematoendothelial population. (A) Summary of RNA-seq comparison between *PF* and *F*. MA plot is shown in **Fig. S4**. (B) RNA-seq TPMs for the 384 ETV2-bound *PF*-gained genes in the *PF* (x-axis) and the *F* (y-axis) populations. (C) Number of upregulated, downregulated, or unchanged genes in *F* vs. *PF* among the 384 ETV2-bound *PF*-gained genes. The upregulated ETV2-bound *PF*-gained genes are termed the “second-wave genes.” (D) Top 10 most over-represented GO terms in the 123 second-wave genes. (E) Relationship between first-wave genes, second-wave genes, and VEGF-induced genes in *PF*. (F) Summary of ATAC-seq comparison between PF and F. (G) Transcription factor (TF) motifs over-represented (odds ratio > 1) within gained ATAC sites in *F* vs. *PF*. (H) Fraction of *F*-vs-*PF* differentially accessible sites intersecting ETV2-binding sites. (I) Relationship between ETV2-bound ATAC sites gained in *F* and ETV2-bound ATAC sites gained in *PF*. (J) RNA-seq TPMs for the 1,090 ETV2-bound *F*-gained genes in the *PF* (x-axis) and the *F* (y-axis) populations. (K) Number of upregulated, downregulated, or unchanged genes in *F* vs. *PF* among the 1,090 ETV2-bound *F*-gained genes. The upregulated ETV2-bound *F*-gained genes are termed the “third-wave genes.” (L) Top 10 most over-represented GO terms in the 244 third-wave genes. (M) TF-encoding second-wave and third-wave genes. Genes are classified by their known roles in endothelial and/or hematopoietic development and function. (N) Summary of the paper. ETV2-bound genes can be classified into the first-wave, second-wave, and third-wave genes, distinguished by the timings of ETV2 binding and transcriptional activation.

### ETV2-binding in the PDGFRα^−^ FLK1^+^ population is associated with activation of the hematoendothelial sublineage gene expression programs

We observed that several well-characterized ETV2 target genes such as *Gata2*, known for hematopoietic specification, and *Tie2*, known for endothelial lineage specification (Koyano-Nakagawa and Garry, 2017) were neither first-wave nor second-wave genes that gained accessibility at ETV2-binding sites in the *PF* population. Therefore, we hypothesized that there might be genes bound by ETV2 *de novo* in the *F* population without prior ETV2 binding in the *PF* population. We therefore investigated the chromatin accessibility landscape in the *F* population by ATAC-seq and compared it with the accessibility landscape of the *PF* population. Of 160,393 union accessible sites in the *PF* and *F* populations, 20,349 sites (13%) gained accessibility and 15,603 sites (9.7%) lost accessibility in the *F* population relative to the *PF* population (**Fig. 4F; Fig. S4D; Table S5**). The sites that gained accessibility in the *F* population were most strongly overrepresented for the Ets family motif (top scoring TF motif, ETV6) (**Fig. 4G; Fig. S4E**) and overlapped with 2,052 ETV2-binding sites with strong statistical overrepresentation (Fisher’s exact test P<2×10^−16^; **Fig. 4H**). ETV2-binding sites were underrepresented among lost accessibility sites (3 sites; P<2×10^−16^). The 2,052 ETV2-bound gained-accessibility sites in the *F* population (“ETV2-bound *F*-gained sites”) included a vast majority (408; 87%) of the 470 ETV2-bound *PF*-gained sites (**Fig. 4I**), suggesting increased utilization of the ETV2-bound *PF*-gained sites in the *F* population. The remaining ETV2-bound *F*-gained sites (1,644 sites) suggested *de novo* ETV2 binding in the *F* population without prior ETV2 binding in the *PF* population.

To define the genes that gained ETV2 binding in the transition from the *PF* to *F* population, we linked the ETV2-bound *F*-gained sites to genes (ETV2-bound *F*-gained sites within the gene body or 100 kb upstream of the TSS). We identified 1,090 “ETV2-bound *F*-gained genes,” linked only to the ETV2-bound *F*-gained sites without ETV2-bound *PF*-gained sites (**Fig. 4I; Table S1**). These sole ETV2-bound *F*-gained genes were highly over-represented for genes upregulated in the *F* population (244 of the 1,090 genes, 22%; Fisher’s exact test P< 2×10^−16^) but not for genes downregulated in the *F* population (63 of the 1,090 genes, 6%; P=0.9) (**Fig. 4J, K**). We termed these 244 upregulated ETV2-bound *F*-gained genes “third-wave” ETV2-bound genes (**Fig. 4K**). The third-wave genes were strongly overrepresented for the GO term “vasculature development” (P=10^−24^) as well as “myeloid cell proliferation” (P=10^−10^) and “endothelial cell proliferation” (P=10^−10^) (**Fig. 4L**). Consistently, the third-wave genes included TFs important for hematopoietic lineage differentiation including *Runx1*, *Gata2*, *Hhex*, *Sox13*, and *Ebf1* (**Fig. 4M**). The third-wave genes also included the endothelial marker genes *Tie2* and *VE-Cadherin.* In addition, the third-wave genes included ETS TFs operating in endothelial and/or hematopoietic sub-lineages downstream of ETV2, such as *Erg*, *Elk3*, *Fev*, and *Ets1* (**Fig. 4M**). Therefore, *de novo* ETV2 binding in the *F* population preferentially marked genes important for the hematopoietic or endothelial sub-lineages derived from the hematoendothelial population (**Fig. 4N**).

## DISCUSSION

ETV2 is a master transcription factor (TF) for the hematoendothelial lineage development (Koyano-Nakagawa and Garry, 2017; Lammerts van Bueren and Black, 2012; Zhao et al., 2017). Overexpression of ETV2 alone is sufficient to transdifferentiate non-endothelial cells into functional endothelial cells (Bersini et al., 2020; Cakir et al., 2019; Lee et al., 2017; Morita et al., 2015; Wang et al., 2020; Zhao et al., 2018). ETV2 is not known to function in cell lineages other than the hematoendothelial lineage (Koyano-Nakagawa and Garry, 2017; Lammerts van Bueren and Black, 2012; Zhao et al., 2017). In this regard, our observation that ETV2 is strongly induced and binds to hematoendothelial genes in the multipotent PDGFRα^+^ FLK1^+^ mesoderm progenitor prior to the hematoendothelial fate commitment is striking. We find that ETV2-dependent gene expression of a large number of ETV2-bound genes occurs upon hematoendothelial fate commitment, not in the common hematoendothelial/cardiac PDGFRα^+^ FLK1^+^ progenitor population, despite ETV2 binding. We propose that ETV2 thereby primes the ETV2-target hematoendothelial genes in the multipotent cardiovascular progenitor prior to hematoendothelial lineage commitment (**Fig. 4N**).

### VEGF is dispensable for mESC-derived cardiomyocyte differentiation

Our time course resolved the sequential order of production of mesodermal progenitors from PDGFRα^−^ FLK1^−^ to PDGFRα^+^ FLK1^−^ (*P*) to PDGFRα^+^ FLK1^+^ (*PF*) to PDGFRα^−^ FLK1^+^ (*F*). We found that the transition from the *PF* multipotent cardiovascular progenitor to the *F* hematoendothelial population requires exogenous VEGF, consistent with the requirement of VEGF or the VEGF receptor FLK1 for hematoendothelial commitment (Kennedy et al., 1997; Shalaby et al., 1995, 1997; Yamashita et al., 2000). However, we find that production of *PF* from the *P* early mesoderm population does not require VEGF, consistent with the previous observation that mESCs deficient for *Vegfa* encoding VEGF-A, are able to differentiate into FLK1-positive cells (Zhao and Choi, 2017). These observations indicate that generation of the *PF* multipotent cardiovascular progenitor is not VEGF dependent. This *PF* population defines the cardiac progenitor and is derived as efficiently with and without VEGF. Furthermore, we found that the *PF* population possesses the same cardiomyogenic potential whether derived with or without VEGF. Thus, VEGF is not required for cardiomyocyte differentiation from mouse embryonic stem cells (mESCs), although VEGF has been used almost universally in mESC-derived cardiomyocyte differentiation protocols (Kattman et al., 2006, 2011; Kokkinopoulos et al., 2016; Mummery et al., 2012). We observed that the *PF* population rapidly transitions to the *F* population in the presence of VEGF and loses the cardiomyogenic potential within 24 hours. We therefore suggest that generation of cardiac progenitors for *in-vitro* analysis should be performed without VEGF.

### Three classes of ETV2-bound genes

We identified three classes of ETV2-bound genes distinguished by the timing of ETV2 binding and transcriptional activation, referred to as the first-wave, second-wave, and third-wave genes (**Fig. 4N**). The first-wave genes were bound by ETV2 in *PF* and upregulated in the *PF* population relative to the *P* population. The first-wave ETV2-bound genes included genes activated in response to VEGF in *PF*. The VEGF-responsive first-wave genes were composed of 7 genes including *Etv2* itself and *Tal1*, a known direct transcriptional target gene of ETV2 (Kataoka et al., 2011; Liu et al., 2015; Wareing et al., 2012). *Tal1* expression immediately follows *Etv2* expression during mESC differentiation (Zhao and Choi, 2017) and requires *Etv2 in vivo* (see accompanying paper by Sinha et al.), supporting the conclusion that ETV2 directly upregulates *Tal1* in *PF*. ETV2 and TAL1 are situated at the top of the hematoendothelial transcriptional hierarchy and negatively regulate the cardiomyogenic fate (Liu et al., 2012; Org et al., 2015; Van Handel et al., 2012). Nonetheless, our data found that the first-wave gene expression does not reduce the alternative cardiomyogenic fate potential of the *PF* population. Thus, the first-wave genes apparently prime, but do not establish commitment of, the hematoendothelial fate in the *PF* population.

Second-wave genes were bound by ETV2 in *PF* but activated in the subsequent *F* hematoendothelial population. These genes represent “poised” loci with ETV2 binding preceding transcriptional activation. These genes include TFs implicated in early vasculogenesis such as *Fli1*, *Sox7*, and *Sox18* (Li et al., 2015; Yao et al., 2019), characterizing the function of hemato-endothelium. Because the level of ETV2 expression plateaus in *PF* and is similar in *PF* versus *F*, hematoendothelial fate commitment in *F* may be regulated by mechanisms that convert poised ETV2 binding to productive transcription at the second-wave genes. A previous study reported that the ETS family transcription factor ETS1 is acetylated by VEGF signaling, and acetylated ETS1 promotes RNA Polymerase II pause release through interaction with a pause release factor BRD4 (Chen et al., 2017). Whether similar post-translational modifications participate in ETV2 target regulation will be an important arena for future investigation.

Third-wave genes were bound by ETV2 in *F* and activated in *F*, demonstrating delayed ETV2 binding compared to the first- and second-wave genes. The third-wave genes include TFs implicated in lineage decisions downstream of the committed hemato-endothelium, such as hematopoietic TF genes *Ebf1*, *Runx1*, and *Gata2*, endothelial TF genes *Erg* and *Ets1*, and other endothelial genes including *VE-cadherin* and *Tie2*. Therefore, ETV2-dependent specification of the hematopoietic and endothelial sub-lineages within the hematoendothelial progenitor must be regulated by ETV2 binding to the third-wave gene enhancers following commitment of the hematoendothelial lineage. The mechanism underlying distinct selectivity of ETV2 binding at second- and third-wave loci is not currently understood. However, ETV2 might promote a hematopoietic gene regulatory network indirectly, for example through *Tal1* activation, as suggested by the accompanying paper by Sinha et al., and consistent with the previous report that *Tal1* can rescue the hematopoietic defects resulting from *Etv2* deficiency (Wareing et al., 2012). Collectively, our data present a model for how ETV2 might control specific stages of hematoendothelial development including in order hematoendothelial fate priming, commitment, and sub-lineage specification.

### Hematoendothelial fate priming by ETV2

Our data presents both similarities and differences with the previously described features of cell lineage priming. A well-described molecular form of lineage priming is “multilineage transcriptional priming”, in which lineage-specific genes are co-expressed at low levels in individual multipotent progenitors (Hu et al., 1997; Laslo et al., 2006; Miyamoto et al., 2002). Our observation that first-wave hematoendothelial genes, including *Etv2* and *Tal1*, are expressed in the *PF* population together with cardiomyogenic TFs is consistent with this form of transcriptional priming. However, unlike the general description of transcriptional priming, *Etv2* was among the highest transcribed TF genes in the *PF* population. Nevertheless, the *PF* population with strong *Etv2* expression retains cardiomyogenic potential. Thus, low expression levels do not seem to be a requisite for transcriptional priming. Furthermore, even at high levels of *Etv2* expression, transcriptional priming did not inhibit the development of alternate fates. Another molecular form of lineage priming is “enhancer priming”, in which lineage-specific enhancers are bound by lineage-specific TFs or associated with histone H3 lysine 4 mono- or dimethylation or increased chromatin accessibility in multipotent progenitors prior to transcriptional activation of associated genes (Goode et al., 2016; Gualdi et al., 1996; Kontaraki et al., 2000; Lupien et al., 2008; Rada-Iglesias et al., 2011). Our observation that the second-wave genes gain accessibility at gene-distal ETV2-binding sites in the *PF* population prior to transcriptional activation in the *F* population is consistent with the form of enhancer priming by ETV2. A well-described mechanism for enhancer priming involves pioneer TFs, a group of TFs capable of binding to nucleosomal DNA and exposing regulatory elements by nucleosome eviction or sliding (Iwafuchi-Doi and Zaret, 2016). These pioneer TFs are suggested to act as a placeholder for other structurally-related TFs to access the exposed regulatory elements (Spitz and Furlong, 2012). Considering the fact that a series of ETS family transcription factors were expressed in the *F* population downstream of ETV2, one hypothesis is that ETV2 acts as a pioneer TF to make enhancers accessible and placehold them in the *PF* population for other ETS family TFs. In support of this hypothesis, some ETV2-binding sites have been reported to be occupied by the ETS family TF FLI1 following ETV2 binding (Liu et al., 2015).

### Fate intermediate isolation to study fate transition dynamics

The present study examines fate intermediates originating from a single time point of mESC differentiation, thereby allowing us to investigate the molecular dynamics during a fate transition with a high temporal resolution. This approach contrasts with the conventional approach to study cell differentiation that compares samples from different differentiation time points and does not capture molecular dynamics during fate transitions (Paige et al., 2012; Stergachis et al., 2013). Our approach also presents advantages over single-cell approaches of investigating fate transitions. While single-cell transcriptome and chromatin accessibility profiling have detected gradual transitions of transcriptome and chromatin profiles during cell fate transitions (Bendall et al., 2014; Buenrostro et al., 2018; Ma et al., 2020; Trapnell et al., 2014), these transition dynamics can only be mapped along inferred developmental time scales. Single-cell approaches do not allow direct evaluation of the fate potentials of the cells under investigation. Our approach enables unambiguous determination of temporal relationships between the sorted fate intermediates and evaluation of the fate potential of intermediates. A limitation of our approach is that our data represent averages of transcriptional states, chromatin states, or fate potentials of cells within the fate intermediates. A future investigation combining fate intermediate isolation with single-cell approaches will likely address the potential heterogeneity of cells within fate intermediates. Regardless of this limitation, our approach defined dynamics of transcription, chromatin accessibility, and ETV2 binding during hematoendothelial fate transition, unveiled pervasive enhancer priming by ETV2 in multipotent cardiovascular progenitors prior to hematoendothelial fate commitment, and resolved the ETV2-dependent waves of transcription that drive successive stages of hematoendothelial development.

## ACKNOWLEDGEMENTS

We thank technical assistance from the University of Chicago Functional Genomics Core, the University of Chicago Cytometry and Antibody Technology Core, and the University of Chicago Light Microscopy Core. This work was funded by NIH Grants R01 HL55337 to K.C., R01 DK119621 and P01 HL146366 to B.L.B., R21/R33 AG054770 to K.I. and I.P.M., R01 HL092153, R01 HL124836, and R01 HL126509 to I.P.M; support was also provided by NIH Grants F30 HL131298 for R.D.N., T32 GM007183 for J.D.S. and E.H., T32 HL007381 for J.D.S., C.K., R.D.N. and A.D.H., T32 GM007197 for A.D.H., and an American Heart Association Western States Affiliate postdoctoral fellowship 16POST30740016 for T.S.

## AUTHOR CONTRIBUTIONS

Conceptualization, J.D.S., C.K., T.S., B.L.B., K.I., and I.P.M.; Methodology, C.K. and I.P.M.; Formal Analysis, J.D.S., K.I., Z.W., and I.P.M.; Investigation, C.K., R.D.N., A.D.H., E.H., and J.K.; Data Curation, J.D.S. and K.I.; Writing – Original Draft, J.D.S., K.I., and I.P.M.; Visualization, J.D.S. and K.I.; Supervision, J.M.C., K.I., and I.P.M.; Project Administration, I.P.M.; Funding Acquisition, J.M.C., K.I., and I.P.M.

## SUPPLEMENTAL MOVIE & TABLES

**Supplemental Movie 1**

Representative beating cardiomyocytes derived from the *PF* population at 4 days after isolation.

**Supplemental Table S1**

Differentially expressed genes, ETV2-bound genes, and first/second/third-wave genes.

**Supplemental Table S2**

Genomic coordinates for the differentially accessible sites between *P* and *PF*.

**Supplemental Table S3**

Genomic coordinates for the 3,866 ETV2-binding sites.

**Supplemental Table S4**

Genomic coordinates for the differentially accessible sites between *PF* derived in ABV and *PF* derived in AB.

**Supplemental Table S5**

Genomic coordinates for the differentially accessible sites between *PF* and *F*.

## METHODS

### Mouse embryonic stem cells

We used the ZX1 mouse embryonic stem cell (ESC) line (Iacovino et al., 2011) for transcriptome and chromatin analyses. We used the Bry-GFP mESC line (Fehling et al., 2003) for characterization of mESC differentiation. These two mESC lines were indistinguishable in their pluripotent states and differentiation potentials.

mESCs were maintained in a serum-free and feeder-free culture system (Gadue et al., 2006; Kattman et al., 2011; Ying et al., 2003, 2008). For ESC maintenance, Neurobasal medium (ThermoFisher Scientific Cat. 21103049) and DMEM/F12 (ThermoFisher Scientific Cat. 10565018) (1:1) were supplemented with 0.5x N-2 supplement (ThermoFisher Scientific Cat. 17502048,), 0.5x B27 supplement (ThermoFisher Scientific Cat. 17504044), 1x penicillin-streptomycin (ThermoFisher Scientific Cat. 15140148), 2mM Glutamine (ThermoFisher Scientific Cat. A2916801), 150μM mono-thioglycerol (MilliporeSigma, Cat. M6145), 0.05% BSA (MilliporeSigma, Cat. A9576), 1000 unit Mouse LIF (MilliporeSigma Cat. ESG1106), 1 μM PD0325901 (Procell Cat. 04-0006-10), 3 μM CHIR99021 (Procell Cat. 04-0004-10). PD0325901 and CHIR99021 were removed 2 days before the initiation of differentiation.

### Mesoderm induction

To initiate differentiation, ESCs were dissociated with TrypLE express (ThermoFisher Scientific) and cultured in a 3:1 mixture of IMDM (ThermoFisher Scientific Cat. 12440053) and Ham’s F12 (ThermoFisher Scientific Cat. 11765054) medium supplemented with 0.5x N-2 supplement, 0.5x B27 supplement, 1x penicillin-streptomycin, 2mM Glutamine, 0.5 mM ascorbic acid (MilliporeSigma Cat. A4544), 450 μM mono-thioglycerol, 0.05% BSA at the density of 0.1 million cells per mL in a 10-cm petri dish (Becton Dickenson) for inducing embryoid bodies (Gadue et al., 2006; Kattman et al., 2011). After 48 hours, the embryoid bodies (EBs) were dissociated with TrypLE express. For mesoderm induction, the dissociated EBs were re-aggregated in the ABV regimen, defined as the StemPro-34 SFM medium (ThermoFisher Scientific Cat. 10639011) supplemented with 2mM Glutamine, 0.5 mM ascorbic acid, 450 μM mono-thioglycerol, 200 μg/ml human transferrin (MilliporeSigma Cat. T8158), 6 ng/ml human bFGF (R&D systems Cat. 233FB), 1 ng/ml human BMP4 (R&D systems Cat. 314BP), 8 ng/ml human Activin A (R&D systems Cat. 338AC), 5 ng/ml mouse VEGF (R&D systems Cat. 494MV). For mesoderm induction in the AB regimen, the culture medium without 5 ng/mL VEGF was used. Flk1 inhibitor, 1.2 μM ZD6474 (SelleckChem Cat. S1046) was treated from day 2 to day 3.75.

### Hematopoietic and cardiac lineage induction

For hematopoietic lineage and cardiac lineage induction, mesodermal cells were sorted (see *Flow cytometry* section) and cultured in the StemPro-34 SF medium supplemented with 2 mM Glutamax-I (ThermoFisher Scientific Cat. 35050-061), 1 mM ascorbic acid, and 30 ng/ml bFGF (Kattman et al., 2006, 2011) for indicated duration at the density of 0.2 million cells per ml in the individual wells of a 48-well flat bottom plate (Becton Dickenson) coated with gelatin (MilliporeSigma).

### Flow cytometry

For cell sorting based on PDGFRα and Flk1 expression levels, embryoid bodies under mesoderm induction in the ABV or AB regimen were harvested and dissociated at the indicated time and incubated with the PE-conjugated anti-PDGFRα antibody (ThermoFisher Scientific Cat. 12-1401-81) and the PE/Cy7-conjugated anti-Flk1 antibody (Biolegend Cat. 359911) for 1 hour on ice. Cells were sorted by the PDGFRα and Flk1 staining levels into the StemPro34 medium using the FACSAria II sorting instrument (BD Bioscience).

For cardiac troponin (cTnT) staining, Day 8 cells were harvested and fixed with cytofix/cytoperm fixation/permeabilization solution (BD Bioscience, Cat. 554714) following manufacturer’s instruction. Cells were incubated with BV421-labeled anti-cTnT antibody (BD Bioscience Cat. 565618) for 1.5 hr and analyzed by LSRFortessa cell analyzer (BD Bioscience). Flowjo software was used to visualize flow cytometry data.

### Immunofluorescence

For immunohistochemistry, the Day 8 cells were fixed with 4% formaldehyde (ThermoFisher Scientific) for 24 hr and washed with PBS. Cells were permeabilized in PBS containing 0.25% Triton X-100 for 10 min. The anti-cTnT antibody (Thermo Fisher Scientific, Cat. MS-295) and anti-myosin heavy chain class II (Abcam Cat. ab55152) antibodies were used.

### Blood colony-forming assay

Cells were plated at 2 days after cell sorting (Day 6 post differentiation) into a methylcellulose-based medium containing hematopoietic cytokines (MethoCult M3434, StemCell Technologies, Cat. 03434) at 6000 cells/ml. Hematopoietic colonies were counted by manufacturer’s instructions at 10 days post-plating (Day 16 of differentiation).

### Quantitative RT-PCR

Quantitative real-time RT-PCR was performed using an ABI Prism 7500 Fast SDS (Applied Biosystems). Total RNA was extracted using NucleoSpin RNAII kit (Takara, Cat#740955.50). qRT-PCR was performed on 96-well optical reaction plates with one-step SYBR Green PCR master mix (Bio-Rad Laboratories, Cat#1725150).

### RNA-seq

Library preparation was performed by the University of Chicago Genomics Facility using the Illumina TruSeq RNA Sample prep kit v2 (Part #RS-122-2001). Library fragments were approximately 275bps in length and were quantitated using the Agilent Bio-analyzer 2100 and pooled in equimolar amounts. Single-ended, 51bp sequencing was performed on the Illumina HiSeq2500 in Rapid Run Mode by the University of Chicago Facility.

### RNA-seq data preprocessing

Transcripts were aligned to the indexed reference for mm10 with default settings using TopHat2 v2.1.1 (Kim et al., 2013; Trapnell et al., 2009). Reads were filtered using bamtools using the following settings: -isDuplicate false -mapQuality “>10” (Barnett et al., 2011). Transcript reads (TPMs) were counted post-alignment using StringTie (Pertea et al., 2015, 2016). TPMs are listed in **Table S1**. Raw and processed RNA-seq data are available at GEO with an accession number GSE136692.

### Differentially expressed genes

Differential expression testing was performed using edgeR v3.16.5 (McCarthy et al., 2012; Robinson and Smyth, 2007, 2008; Robinson et al., 2010; Zhou et al., 2014) and limma v3.30.13 (Ritchie et al., 2015) packages in R v3.3.2. Low level genes were removed within each condition using median log_2_-transformed counts per gene per million mapped reads (cpm) of 1 and a union generated from those lists. Differential expression testing was performed using a general linear model (GLM) framework. For comparing FAB and FAB+V, a covariate for replicate was included to correct for batch effect. Genes with absolute log_2_ fold change greater than 0.5 and false discovery rate (FDR) smaller than 5% were defined as differentially expressed genes. The human transcription factor annotation reported in Lambert et al (Lambert et al., 2018) was downloaded from http://humantfs.ccbr.utoronto.ca/download.php and incorporated into the mouse gene list based on the gene symbols. Differentially expressed genes are listed in **Table S1**.

### ATAC-seq

ATAC-seq (Assay for Transposase-Accessible Chromatin using sequencing) was performed as previously described (Buenrostro et al., 2015). Libraries were amplified and normalized with the Illumina Nextera DNA Library prep kit (FC-121–1031) according to the manufacturer’s protocols. Libraries were quantitated using the Agilent Bioanalyzer, pooled in equimolar amounts, and sequenced with 50-bp single-end reads on the Illumina HiSeq following the manufacturer’s protocols through the Genomics Core Facility at the University of Chicago.

### ATAC-seq data preprocessing

Sequencing reads were aligned to the mm10 genome using Bowtie2 v2.3.0 (Langmead and Salzberg, 2012) and SAMtools v0.1.19 (Li, 2011; Li et al., 2009). Peak calling was performed using MACS2 callpeak (Feng et al., 2012; Zhang et al., 2008) using the settings --nomodel --shift −100 --extsize 200 -q 0.1 after pooling biological replicates. A fold-enrichment track was generated using MACS2 using the bdgcmp function (-m FE) for visualization on the genome browser. Following removal of ENCODE blacklist sites (Amemiya et al., 2019; ENCODE Project Consortium, 2012), a union set of sites was generated by identifying summits that overlapped within 200bp and arbitrarily selecting the summit of highest positional value. Summits were then extended by 200bp in both directions to create the set of union sites. The union set of sites utilizes biological groups not explicitly described in this manuscript, but are relevant and publically accessible through GEO. Fold-enrichment scores were assigned to each site using the multiBigwigSummary function from deepTools (Ramírez et al., 2016).

### Differentially accessible sites

For differentially accessible site analysis, we first extracted ATAC sites that were identified in either of the two datasets under comparison or boh. The fold-enrichment scores for this set of ATAC sites were processed using a general linear model in edgeR v3.16.5 (McCarthy et al., 2012; Robinson and Smyth, 2007, 2008; Robinson et al., 2010; Zhou et al., 2014) and limma v3.30.13 (Ritchie et al., 2015) packages in R v3.3.2 to identify differentially accessible sites. Differentially accessible sites with the P-value less than 10^−3^ and absolute log_2_ fold change greater than 1 were selected for downstream analyses. The list of differentially accessible sites are listed in **Table S2** (*PF* vs. *P*), **Table S4** (*PF* derived in ABV vs. *PF* derived in AB), and **Table S5** (*F* vs. *PF*).

### ETV2-binding sites

We used the previously published ETV2 ChIP-seq data performed in a FLK-positive cell population derived from mESC differentiation (Liu et al., 2015), available at GEO under the accession ID GSE59402. This dataset used a mESC cell line that carries a V5-epitope tagged ETV2 transgene under the doxycycline-inducible promoter (Liu et al., 2015). In the experiment, the V5-epitope tagged ETV2 was expressed from Day 2 to Day 3.5 of differentiation by doxycycline, and ETV-associated chromatin was immunoprecipitated using an anti-ETV2 antibody or an anti-V5 tag antibody (Liu et al., 2015). We aligned sequencing reads to the mm10 mouse reference genome using Bowtie2 (Langmead and Salzberg, 2012) with the default “--sensitive” parameter. Reads with MAPQ scores greater than 20 were used in downstream analyses. Reads from biological replicates of ChIP and the corresponding input were processed by MACS2 (v2.1.0) (Zhang et al., 2008). Aligned reads from three ChIP-seq replicates (one replicate from anti-ETV2 ChIP and two replicates from anti-V5 ChIP; Short Read Archive IDs SRR1514692, SRR1514695, SRR1514696) and aligned reads from two control ChIP experiments (one replicate from non-ETV2 induced cells and one replicate from IgG control; Short Read Archive IDs SRR1514691 and SRR1514694) were used in MACS2 program (Feng et al., 2012; Zhang et al., 2008) to identify statistically overrepresented peak regions and peak summits and to produce fold-enrichment scores. The parameters used in MACS2 were [callpeak -g hg --nomodel --extsize 200 --call-summits]. Identified ETV2-enriched regions with the p-value < 10^−10^ and the fold enrichment score greater than 4 that did not overlap ENCODE mm10 blacklisted regions (https://www.encodeproject.org/annotations/ENCSR636HFF/) or the mitochondrial genome were selected. This yielded 3,868 ETV2-binding sites (**Table S3**). An ATAC site was considered possessing ETV2-binding sites when the 400-bp region centered around the ATAC site summit overlapped the summit of at least one ETV2-binding site.

### Gene ontology analysis

Gene ontology (GO) analyses were performed using Metascape (Tripathi et al., 2015). Gene Symbols of the genes of interest were used to examine overrepresentation of Biological Process and Molecular Function GO terms with default parameters (minimum gene count 3, P <0.01, enrichment over background >1.5). P-values were derived from cumulative hypergeometric statistical tests and computed in Metascape (Zhou et al., 2019). Reported GO terms in the figures are the “Summary” GO terms of all associated GO terms, and the number of genes represent the union of genes affiliated with the associated GO terms.

### DNA motif analysis

We used FIMO in the MEME suite (v5.0.5) (Grant et al., 2011) to scan for the presence of TF motifs in each ATAC site (+/–100 bp from ATAC summit). We used the *Mus Musculus* CIS-BP TF motif database (v2.0.0) (Weirauch et al., 2014), which contains Position Weight Matrix and TF family annotation for each TF. For each ATAC site and for each TF, we determined whether at least one motif is present or not. We then performed, for each TF motif, Fisher’s exact test with a contingency table with counts for ATAC sites with motif presence or not and differentially accessible or not. This resulted in, for each motif for each ATAC group, a P-value, odds ratio, and motif-containing fraction of ATAC sites. To find overrepresented motifs, we selected motifs with odds ratio greater than 1 and the 95% confidence interval of the odds ratio not overlapping 1. We then selected a TF motif with the highest score defined by log_2_(odds ratio) x fraction for each motif family. The TF name, TF family name, fraction, and the P-value are plotted in figures.

### Association between ATAC sites and genes

One ATAC site was linked to one gene when the ATAC site summit resided in the gene body or 100 kb upstream of the transcription start site (TSS) of that gene. ATAC sites that did not fulfill this condition were not linked to genes. We used the following algorithm to select a single gene when multiple genes could be linked: a) ATAC sites located in the gene body of one gene and within 100 kb upstream of another gene were assigned to the gene whose gene body contains the ATAC sites; b) ATAC sites located in the gene body of two different genes were assigned to the gene whose TSS was closer to the ATAC sites; and c) ATAC sites located within 100 kb upstream of two different genes were assigned to the gene whose TSS was closer to the ATAC sites.

### Single-cell RNA-seq analysis

Single-cell RNA-seq data was downloaded from GEO (GSE130146) (Zhao and Choi, 2019) and imported into R using Seurat package version 4.0.0. In preprocessing, genes expressed in at least three cells were kept, and cells with less than 5% mitochondria read and greater than 2000 unique genes were kept. This preprocessing resulted in 16,249 unique genes and 2,202 cells. Genes with read counts greater than 0 were considered expressed in a cell, and genes with read counts equal to 0 were considered not expressed in a cell. To obtain expression levels, read counts were first normalized within each cell by dividing by the total counts of each cell, then multiplied by 10000, and then natural-log transformed, following the “LogNormalization” method in the Seurat package. The single-cell *P* population was defined as *Pdgfra*-expressed, *Kdr*-not-expressed cells; the single-cell *PF* population was defined as *Pdgfra*-expressed, *Kdr*-expressed cells; and the single-cell *F* population was defined as *Kdr*-expressed, *Pdgfra*- not-expressed cells. To compute statistical significance of *Etv2*-expressed cell count differences between *P* and *PF*, Fisher’s exact test examined the association between *Kdr*-expressed cells and *Etv2*-expressed cells within *Pdgfra*-expressed cells. For the comparison between *PF* and *F*, Fisher’s exact test examined the association between *Pdgfra*-expressed cells and *Etv2*-expressed cells within *Kdr*-expressed cells. Wilcox rank-sum test examined the statistical significance of *Etv2* expression level differences between *P* and *PF* and between *PF* and *F*, using normalized read counts.

### Reference genome

We used mouse reference genome mm10 for all data analyses.

### Data availability

RNA-seq and ATAC-seq data are available at GEO with an accession number GSE136692.

---

**Figure S1.**
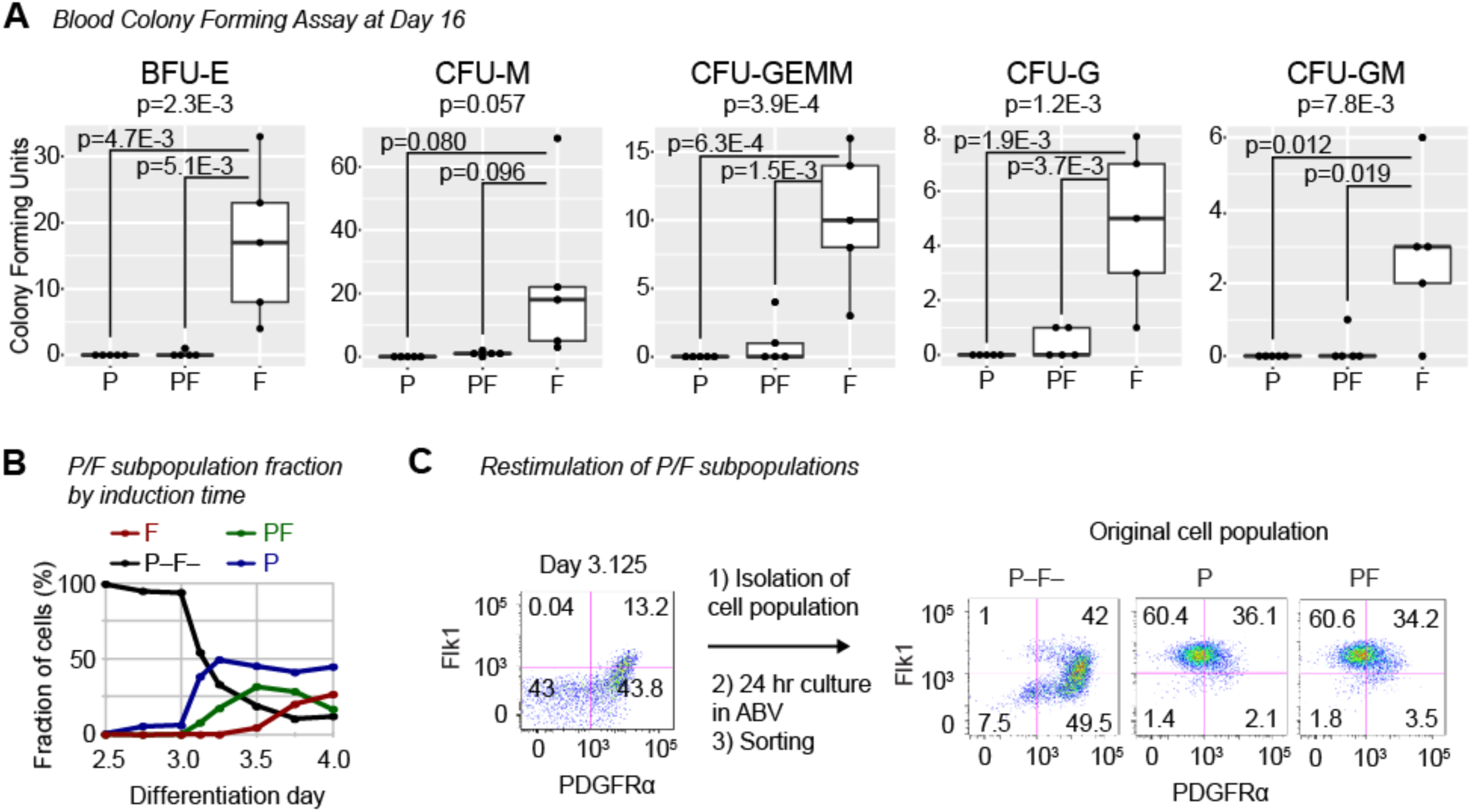
Characterization of PDGFRα/FKL1 subpopulations (related to Figure 1) (A) Blood colony forming assays for individual blood subtypes. BFU-E, erythroid burst-forming unit. CFU-M, megakaryocyte forming unit. CFU-GEMM, granulocyte, erythrocyte, monocyte, megakaryocyte-forming unit. CFU-G, granulocyte-forming unit. CFU-GM, granulocyte, monocyte-forming unit. (B) PDGFRα/FKL1 subpopulation composition during differentiation. FACS plots are shown in main Figure 1D. (C) Restimulation of isolated PDGFRα/FKL1 subpopulations in the ABV regimen for 24 hours.

**Figure S2.**
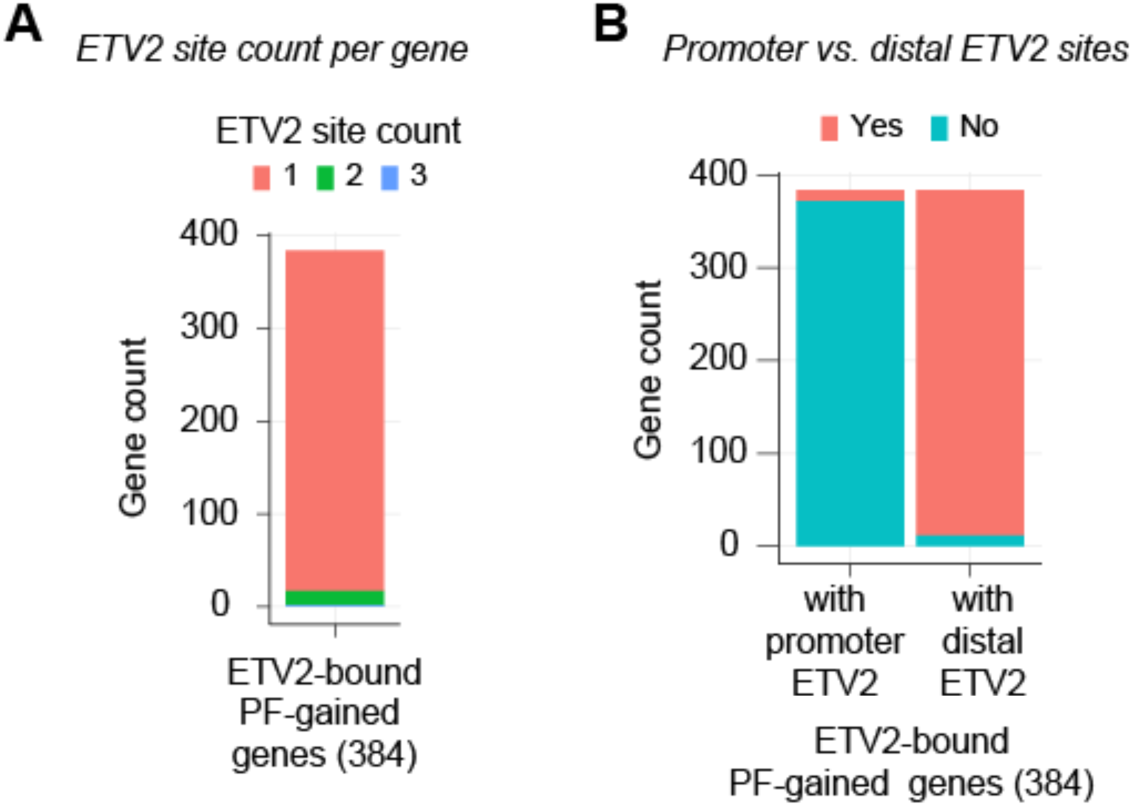
ETV2-bound PF-accessibility gained genes (related to Figure 2) (A) ETV2-bound *PF*-ATAC gained genes are grouped by the number of linked ETV2-bound *PF*-ATAC gained sites. (B) Number ETV2-bound *PF*-gained genes linked to ETV2-bound *PF*-gained sites at promoter regions (+/–2 kb of TSS) or distal regions (outside +/–2 kb of TSS).

**Figure S3.**
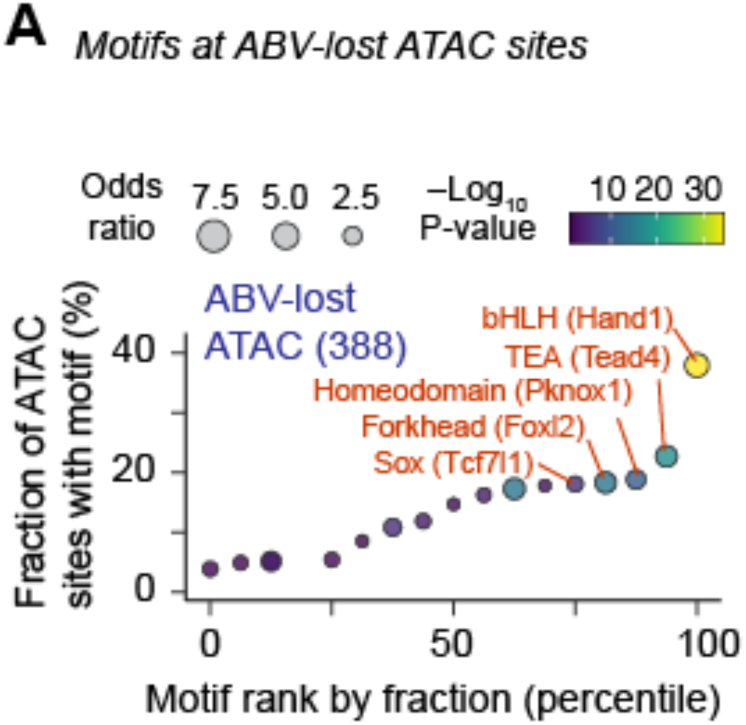
DNA motifs overrepresented in the lost accessibility sites in *PF* derived with ABV vs. *PF* derived with AB (related to Figure 3).

**Figure S4.**
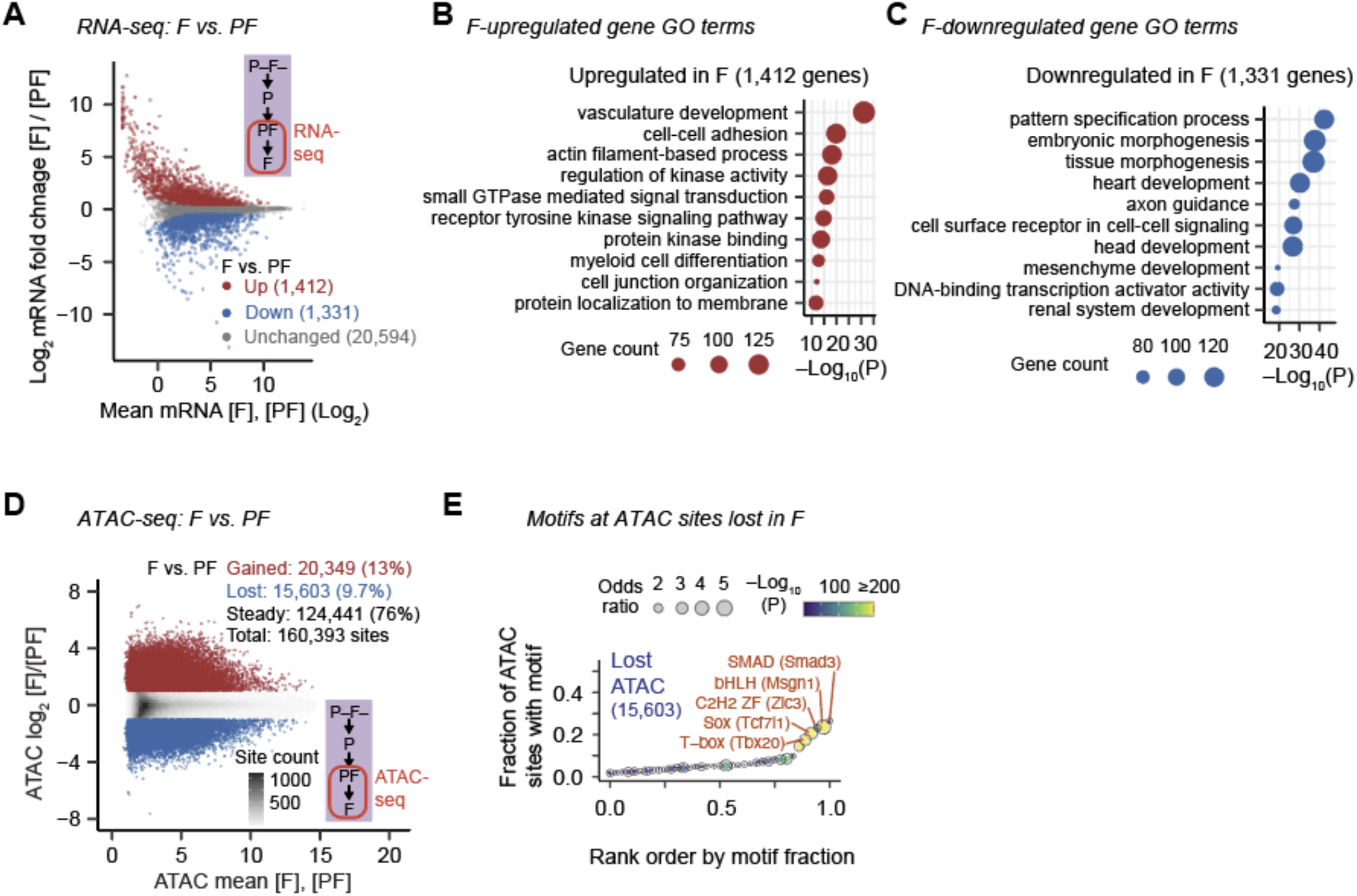
Transcriptional and chromatin accessibility comparison between F and PF (related to Figure 4) (A) MA plot comparing RNA-seq TPMs (transcripts per million) of all genes in *F* vs. *PF* populations. (B) Top 10 most over-represented GO terms in the upregulated genes in *F* vs. *PF*. (C) Same as (B), but GO terms for the downregulated genes are shown. (D) MA plot comparing ATAC-seq-derived chromatin accessibility at the 160,393 union ATAC sites in the *F* population vs. the *PF* population. (E) Transcription factor (TF) motifs over-represented (odds ratio > 1) within lost ATAC sites in *F* vs. *PF*.

